# How thermostable direct haemolysin (TDH) diverges from TDH-related haemolysin (TRH)? Reassessing haemolysins in *Vibrio parahaemolyticus* through functional and structural representation learning

**DOI:** 10.64898/2026.09.02.748974

**Authors:** Zhuosheng Liu, Yi Zhou, Chengchu Liu, C Titus Brown, Luxin Wang

## Abstract

*Vibrio parahaemolyticus* (*Vp*) is the major foodborne pathogen transmitted via shellfish products, which has posed significant threats to modern public health and resulted in significant economic damage to the seafood industry. Numerous studies have documented diverse aspects of *Vp* pathogenicity, among which thermostable direct haemolysin (TDH) and TDH-related haemolysin (TRH) are considered as the major virulence biomarkers of *Vp*. Despite their established roles as key biomarkers, a systematic understanding of the divergence of TDH and TRH across sequence, structure, and function remains limited. In this study, a multi-scale analysis of TDH and TRH was performed using publicly available (2131 and 99 records from NCBI and Uniprot database, respectively) amino acid sequence data combined with representation learning and structure prediction. Global alignment of curated TDH and TRH sequences revealed extensive, distributed mutations and clear separation between TDH and TRH at the amino acid level (percentage identity of between TDH and TRH ranging from 56.1–67.4%). In contrast, protein language model–derived embeddings showed high global functional similarity while preserving distinct clustering patterns, indicating conserved core functionality alongside nuanced divergence echoed with the structural inference by AlphaFold. Importantly, modeling of mutation trajectories demonstrated that the transition from TDH to TRH is driven by accumulated, genome-wide residue changes rather than a small set of key mutations. Together, these results suggested that TDH and TRH represent functionally conserved yet evolutionarily diverged toxins driven by accumulated sequence variation throughout the full-length amino acid sequence. These accumulated point mutations lead to major structural difference: TRH forms an α-helical tail that TDH lacks, which suggests that TDH and TRH disrupt host membranes by different mechanisms despite their conserved core function such as pore-forming and ion flux induction capability. These methods provided unprecedented detailed insights into the functional and structural properties of *Vp* haemolysin, offering critical information on how multi-dimensional variations in sequence might influence their role in *Vp* pathogenicity. Insights from this study reinforce the rationale for using *Vp* strains harboring *tdh* and *trh* genes in experimental design for environmental fitness investigation.

## Introduction

*Vibrio parahaemolyticus* (*Vp*) is recognized as the leading cause of human gastroenteritis associated with seafood consumption in the United States and worldwide, which serves as an important seafood-borne pathogen and poses significant threats to modern public health (Z. Liu et al., 2024). With extensive records of negative impacts both in public health and economic loss estimated by the CDC and USDA at 45,000 annual illnesses and $40 million in U.S., characterizing its virulence is of critical importance to enhance our understanding and facilitate the development of effective prevention and detection methods (CDC, 2023; USDA, 2013). Over the past several decades, different virulent factors associated with *Vp* have been characterized using different precisely targeted genetics and omics-oriented biotechniques: major virulence factors including thermostable direct haemolysin (TDH), TDH-related haemolysin (TRH), the adhesion molecule MAM7, and two Type III Secretion Systems (T3SS1 and T3SS2), which contribute to the bacterium’s cytotoxicity, enterotoxicity, and overall pathogenicity. Zhang and Orth (2013) grouped these virulent factors into several broad functions including to facilitate attachment, disrupt cellular processes, and invade host cells. Among distinct virulent factors, prior studies have consistently identified TDH and TDH-related haemolysin (TRH) as primary virulence factors of *Vp* (pore-forming toxins disrupting integrity of host cell membrane and causing cell lysis). Both haemolysins play critical roles in mediating cytotoxicity and enterotoxicity during infection and act as the main-driver of gastroenteritis and diarrhea, the common symptoms of Vibriosis (Ceccarelli et al., 2013; Hiyoshi et al., 2010; Z. Liu et al., 2026; Z. Liu et al., 2023; Zhang & Orth, 2013).

However, the comprehensive characterization of major haemolysin of *Vp* in the context of genetic evolution, protein structure, and functional divergence has remained limited currently. Noticeable difference in survival rates and stronger growth dynamics of *Vp* strains harboring *tdh* and *trh* genes were reported in the previous investigation (B. Liu et al., 2016). Z. Liu et al. (2023) reported that a *tdh+ Vp* strain exhibited significantly stronger growth and activation of biosynthetic and energy metabolism pathways than the *trh+* strain at 30 °C, suggesting the potential involvement of TDH- and TRH-associated lineages in *Vp* growth. Ellett et al. (2022) hypothesized the importance of energy costs on virulence factor affecting *Vp* survival and growth outcomes. Given these metabolic and survival/growth trade-offs, these evidence led to an imperative need to reveal how TDH and TRH diverge across amino acid sequence, structure, and function.

With the advancement in next-generation sequencing technology, data-driven omics research has been widely conducted on *Vp* in the past years (Z. Liu et al., 2025). Given these developments, Z. Liu et al. (2024) described the current progress in understanding of *Vp* driven by the advancement of omics-oriented biotechniques over past decades and underscored the importance of integrating genomic, transcriptomic, and proteomic data to elucidate the virulence mechanisms of *Vp*. In this paper, with more than 2,000 complete annotated genomes of *Vp* are available at NCBI, the authors utilized different machine learning algorithms to predict the virulence of *Vp* based on its pangenome and successfully traced back important genes essentially contributing the virulence of *Vp*. Large-scale omics-data and advanced analytical methods powered by diverse learning algorithms demonstrated their potential to complement current understanding of the virulence associated with foodborne pathogens.

The advancement in deep learning has led to representation learning as the cornerstone in biological science especially in the context of amino acid investigation, which is an approach that automatically transforms raw, high-dimensional data into low-dimensional, dense vectors (embeddings) serving as an novel emerging method to decode complex patterns in biological sequences (Iuchi et al., 2021). Among different representation learning methods, the current progress of state-of-the-art (SOTA) protein language models provide a novel and groundbreaking solution into structural biology. For instances, AlphaFold leverages massive multiple sequence alignments (MSAs) through its inside adapted transformer module (Evoformer) to construct high-dimensional latent representations, which then can be used in downstream protein structure predictions and functional inferences across the protein universe (Jumper et al., 2021). In addition, Evolutionary Scale Modeling (ESM) can directly treat amino acid sequences as text, and further utilize transformer-based protein language models to learn evolutionary constraints and biophysical properties via unsupervised pretraining (Lin et al., 2023). SOTA foundation models in molecular biology have revolutionized the field by providing highly accurate structural predictions, enabling unprecedented insights into complex biological data and facilitating the development of predictive tools for understanding protein functions.

In summary, a comprehensive analysis of major haemolysin TDH and TRH is fundamentally critical and is imperatively needed at this moment. To fulfill this knowledge gap, systematic bioinformatics complemented with SOTA pretrain protein language model and protein structure prediction models could provide an unprecedented opportunity to reveal genetic divergence, accurately predict protein structures and elucidate functional mechanisms.

## Materials and Methods

### Amino acid sequence collection and preprocessing

An overview of data collection and downstream data wrangling was illustrated in **S Fig 1**. All available *Vp* haemolysin amino acid sequence (TDH and TRH) were retrieved from NCBI and Uniprot database. Haemolysin of *Vp* amino acid sequence (TDH and TRH) were retrieved from the NCBI Protein database using Biopython’s Entrez and SeqIO modules (Biopython, RRID:SCR_007173). Searching criteria were defined as following querystrings ‘tdh[Gene] OR "thermostable direct haemolysin"[All Fields] OR "thermostable direct haemolysin"[All Fields]) OR (trh[Gene] OR "TDH-related haemolysin"[All Fields] OR "thermostable direct haemolysin- related"[All Fields]) AND "*Vibrio parahaemolyticus*"[Organism]’. A total of 2131 records were hit and retrieved using these criteria from NCBI Genebank. All 2131 protein accessions were downloaded via Entrez.efetch (rettype = “fasta”, retmode = “text”) and written back to a FASTA file. *Vp* haemolysin amino acid sequence (TDH and TRH) were retrieved from UniProt using Biopython (RRID:SCR_007173) with UniProtKB REST API (https://rest.uniprot.org/uniprotkb/search) with following query strings ‘(gene: tdh) AND organism_id:670’, ‘(gene:tdh2) AND organism_id:670’, ‘(gene:tdh3) AND organism_id:670’, ‘(gene:tdh4) AND organism_id:670’, ‘(gene:trh) AND organism_id:670’, ‘(gene:trh2) AND organism_id:670’. A total of 99 records were hit and retrieved using these criteria from Uniprot database.

All 2230 retrieved potential candidates of *Vp* haemolysin amino acid sequence (TDH and TRH) records were further preprocessed. A description of each entry was manually checked to guarantee the correctness of the amino acid sequence being considered a candidate for TDH/TRH. Based on information reported in previous documented studies, only full-length toxins of 189 or 190 amino acids were determined as the sequence length for complete sequence (Nishibuchi & Kaper, 1995). In total, 106 unique complete TDH/TRH protein amino acid sequence were retrieved, of which the labeling of TDH and TRH were manually determined by the description of data entry metadata. Both complete and incomplete sequences were subjected to global pairwise alignment to systematically identify point mutations and residue-level variations across the dataset. Furthermore, complete sequences were embedded using the Evolutionary Scale Modeling (ESM) protein language model to capture high-dimensional contextual representations of amino acid patterns. Besides investigation on functional discrepancy of *Vp* haemolysin, protein structure was inferred by AlphaFold.

### Bioinformatics analytics

With preprocessed TRH and TDH amino acid sequence retrieved from GeneBank and Uniprot database, similarity matrix was constructed using the similarity score after pair-wise global alignment using Clustal Omega in Biopython (Edgar, 2004). The resulting pair-wise alignment scores facilitate the identification of relationships and functional similarities among the studied sequences, ultimately contributing to a deeper understanding of their biological roles. Pairwise sequence similarity was then converted to a distance matrix and hierarchical clustering was performed with average linkage using SciPy’s linkage, where clusters were formed based on the average distance between all pairs of sequences in two groups. The resulting dendrogram clustering showed the hierarchical relationships among TDH/TRH sequences, where branch lengths indicate degree of dissimilarity and cluster groupings indicate relative similarity based on the distance matrix. To further utilize incomplete sequences, multiple sequence alignments (MSAs) were conducted on the identified clusters to refine the comparison and elucidate conserved regions and functional motifs using Clustal Omega (Sievers & Higgins, 2014). The complete TDH sequence with highest per-residue frequency across all alignments was selected as the reference sequence. Position-specific amino acid substitution frequencies were calculated by comparing each aligned sequence to the reference sequence (gap positions and residue on reference sequence were excluded).

For each position, an amino acid accumulation matrix was calculated and then visualized using heatmap highlighting substitution patterns across all TDH/TRH amino acid sequences using both complete only and all sequences (incomplete and complete). The mutation per position was further quantified using amino acid accumulation matrix. A position-mapping process was conducted by scanning non-gap residue between the reference sequence and each of the other amino acid sequence in global alignment results. Afterward, a smoothed mutation trajectory was generated using a centered rolling mean (window size = 5), which aimed to catch the pattern of highly variable and relatively conserved regions in *Vp* haemolysin amino acid sequence.

### Functional representation learning of haemolysin using ESM2

Complete TRH and TDH amino acid sequence were used as input into the pretrained model facebook/esm2_t6_8M_UR50D, which at the end generated embedding vector with dimension size fixed to 320 as the final output on Nvidia DGX Spark Blackwell platform (Lin et al., 2023). Once the representation vectors are obtained, UMAP clustering (cosine similarity used as distance metrics) was selected to preserve the global functional variance of TDH/TRH embedding vectors (McInnes et al., 2018). performed to study the spatial organization of these molecules. This clustering analysis leverage functional similarity from ESM, which allowed for the comparison of TDH/TRH haemolysin in the context of toxin activity. To further investigate the functional divergence of TDH/TRH, the description of amino acid sequence (TDH vs TRH) was treated as distinct functional classes. These complete TDH and TRH amino acid sequence were then one-hot encoding transformed and fit into random forest classifier to predict their functional classes (80% sequences were used as train set whereas 20% sequences was used as test set with five-fold cross validation). To identify the important position contributing to the differentiation of TDH/TRH, importance analysis of trained random forest classifier was performed. The mean points of ESM embedding of TDH and TRH were calculated as the center points and the closest embedded TDH/TRH vectors to the center points were selected as the start points, which served as the foundation of point mutation. The trajectory path from TDH toward TRH was further constructed: the TDH sequence closest to the mean ESM2 embedding of all TDH sequences based on Euclidean distance was used as the starting point. Point substitutions with non-zero feature importance were identified by gradient-boosting trees (multiple random seeds), of which yielded sequences were re-embedded using ESM2. The final trajectory was made new embeddings moving monotonically closer to the mean ESM2 embedding of all TRH sequences center, which consists of 23-steps in total. The set of TDH and TRH ESM2 embedding was the clustered using K-means to locate the embedding serving as the centroid following visualized using UMAP.

### Structural folding representation of *Vp* haemolysin using AlphaFold and downstream simulations

By examining the spatial relationships among the *Vp* haemolysin, the goal of this part aims to elucidate the protein folding characteristics that may influence the biological activity of TDH and TRH. AlphaFold is a deep learning-based model specifically designed for predicting protein folding, which used to be achieved by costly and time-consuming methods such as X-ray crystallography, nuclear magnetic resonance (NMR) spectroscopy, and cryo-electron microscopy (Abramson et al., 2024; Jumper et al., 2021). The folding structure of TRH and TDH were constructed based on AlphaFold2 inference results using ColabFold (Mirdita et al., 2022).

Complete TDH and TRH sequence were parsed into single and independent .fasta files. For each complete sequence, multiple sequence alignments (MSAs) were generated using the UniRef+Environmental search mode against the UniRef30 (2023_02) and PDB100 (230517) databases. After MSAs, five independent models (each model conducted three recycling iterations to improve structural accuracy) were used for protein structure inference using AlphaFold2 and then inferred protein structures were subsequently ranked based on confidence scores (high to low). For each complete sequence, only the predicted structure (.pdb) with highest confidence score was selected for downstream analysis. High-resolution structural visualization and comparative analysis were conducted using PyMOL Version 3.18 (Schrödinger, Inc., software available at pymol.org). Thermodynamics of complete sequence folding structure (.pdb) were evaluated using PyRosetta using the ref2015 score function with fast_relax to obtain each Rosetta Energy Unit (REU). For membrane-affinity evaluation, each Martini coarse-grained model of folding structure was obtained through martinize2 (Wassenaar et al., 2015), which was then placed into a mixed lipid membrane (Phosphatidylcholine: Phosphatidylethanolamine= 70:30) using insane (Wassenaar et al., 2015). Each system was simulated in GROMACS using a standard four-step workflow: energy minimization (EM), temperature equilibration (NVT), pressure equilibration (NPT), and a final production run by default (Wassenaar et al., 2015).

## Results

### Compilation of existing *Vp* haemolysin amino acid sequence from public database

*Vp* haemolysin amino acid sequences were systematically obtained from public databases including NCBI GenBank and UniProt to build the foundation for later characterization of the structural and functional diversity of *Vp* haemolysin. A total of 2,131 records from GenBank and 99 records from UniProt were initially collected respectively according to the search criteria. The downstream filtering based on amino acid annotation and sequence length with deduplication led to a final 342 unique sequences (complete + incomplete) with confirmed identity as TDH/TRH, among which 106 were complete sequences and 236 were partial sequences. Sequence length distribution revealed a clear distinction between complete and incomplete amino acid sequences. It is not uncommon that two haemolysin sequence was labeled as both TDH and TRH in different records (same unique sequence mapping to TDH/TRH across records). These identification-unclear amino acid sequences were labeled as unknown first (**Fig 1a)**. These obtained amino acid-associated strain sources are diverse as shown in **Fig 1.b.** Both TDH and TRH sequences exhibit diversity in length, as evidenced by the spread in the histogram. Based on collected amino acid sequences from public databases, a substantial amount of both TDH and TRH fall below the expected full-length size (∼165–190 amino acids for mature haemolysin), which served as clear evidence that partial or fragmented sequence entries have been widely deposited in public databases. These truncated sequences could be due to incomplete genome assemblies and partial low quality gene sequence lacking start/stop codons. Strain information such as isolation source was noted during data collection revealing diverse isolation source types. Based on the isolation source information, the isolation sources were classified into four categories including clinical, seafood, environmental, and other (**Fig 1b**). Based on existed isolation source information, clinical isolates dominate the dataset, in which stool and patient with gastroenteritis were the top two isolation source. Seafood isolates are the second most common, with oyster the leading isolation source following by fish, clams, and shrimp.

**Fig 1:**
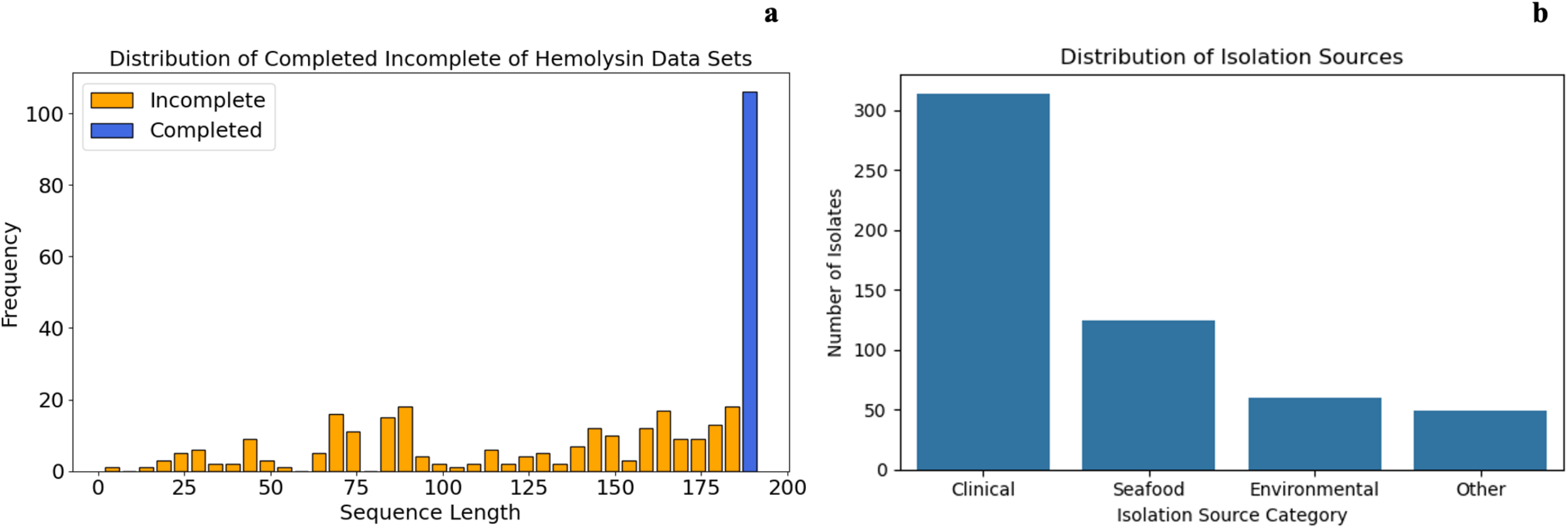
Distribution of sequence lengths for haemolysin sequences of *Vp* collected from public databases (**a**); Distribution of *Vp* strains containing TDH/TRH from public databases (**b**).

Environmental isolates were predominantly from water-related sources such as water, seawater, and mud accounting for the majority. These collected isolation source information of *Vp* harboring TDH/TRH resonated with patterns in previous studies (Z. Liu et al., 2024; Su & Liu, 2007).

### Sequence conservation pattern has been evidenced by global alignments using complete *Vp* haemolysin amino acid sequence

Global alignment enabled a position-resolved quantitative analysis of mutation accumulation using both complete-only and all (complete + incomplete) of *Vp* haemolysin amino acid sequences. Using the aligned reference sequence as an anchor, an amino acid accumulation was constructed to capture substitution frequencies at each position relative to the one of reference sequence. Dense signals of substitution across reference sequence were illustrated in **S Fig 3a**, and the mutation landscape appeared saturated as all available *Vp* haemolysin sequence were included as shown in **S Fig 3b**. This pattern might be due to the alignment noise introduced by incomplete sequences and gap-rich regions, which inflate apparent variability and obscure biologically meaningful signals. In contrast, complete haemolysin sequences improved signal-to-noise resolution and revealed the conservation pattern in mutation landscape. The heatmap revealed sparse and localized substitution patterns (**Fig 2a.)**. Correspondingly, the mutation landscape showed clearer fluctuations including distinct peaks and valleys that were further clarified by rolling mean smoothing (**Fig 2b.)**. Three main conserved regions were found: positions 25-44, 62-122, and 147-182. Yanagihara et al. (2010) reported the N-terminal conserved region in TDH as ‘FELPSVPFPAP’, which was parsed against 106 complete sequences. The regions that match with ‘FELPSVPFPAP’ commonly start at position 25 and end at position 35. The mutation count of this conserved region after smoothing was used as the criteria to discover the remaining conserved regions (**Fig 2**). Regions from position 62 to 122 showed conserved regions of TDH/TRH, in which 31 of 61 positions (50.9%) were invariant, and regions from position 147 to 182 showed conserved regions of TDH/TRH, in which 15 of 36 positions (41.7%) were invariant across all 106 complete sequences.

**Fig 2.**
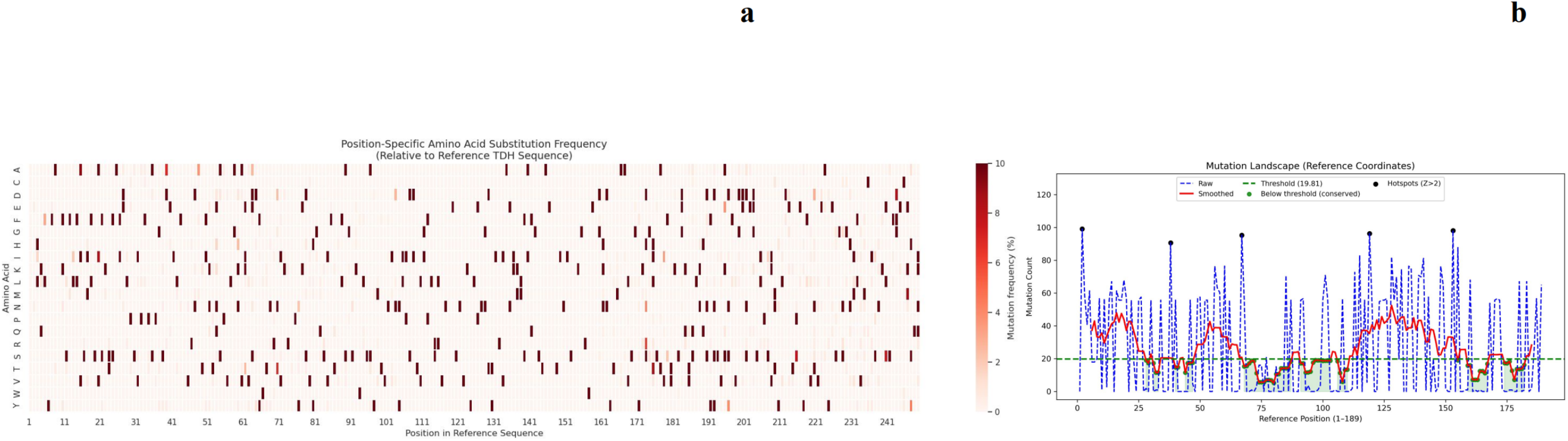
Comparative analysis of position-specific mutation frequencies against reference sequence using aligned complete of *Vp* haemolysin amino acid sequence (a) and mutation landscape of *Vp* haemolysin reference positions (1–189). Raw counts (blue dashed) and smoothed trend (red) are shown; the green dashed line indicates the threshold of conservation with continuous positions below the threshold as conserved (green) and hotspots (Z > 2) marked in black dot (b).

### Distinct divergence has been detected by pair-wise global alignment based on complete *Vp* haemolysin amino acid sequence

To further investigate divergence of *Vp* haemolysin, pair-wise global alignments were conducted. Complete *Vp* haemolysin amino acid sequences with rough labels were then clustered. Based on the results, the similarity matrix provides valuable insights into the relationships between TDH and TRH, in which the color gradients effectively highlight the distribution of similarity scores and two well-separated protein groups with high intra-group similarity and low inter-group similarity were observed (**S Fig 4**.). Hierarchical clustering revealed distinct groupings of haemolysin with initial gathered labeled (**Fig 3**.): hierarchical clustering clearly separated sequences labeled as TDH and TRH into two distinct, well-defined clades, indicating strong divergence between the two groups at the sequence level. Based on dendrogram clustering, three haemolysin sequences exhibiting discordance between their annotated labels and local neighborhood composition were reassigned: seq_76 (TDH/TRH unclear → TDH), seq_30 (TDH → TRH), and seq_70 (TDH/TRH unclear → TRH). Cross-group cosine similarity analysis based on pair-wise global aligned sequences revealed the closest TDH–TRH pair (seq_73 TDH and seq_41 TRH; percent identity 59.69%) and extreme divergence between the most distant pair (seq_72_TDH and seq_60_TRH; percent identity 67.37%), which highlighted substantial sequence-level separation between the TDH/TRH toxins at the amino acid level (**S Fig 5**.).

**Fig 3:**
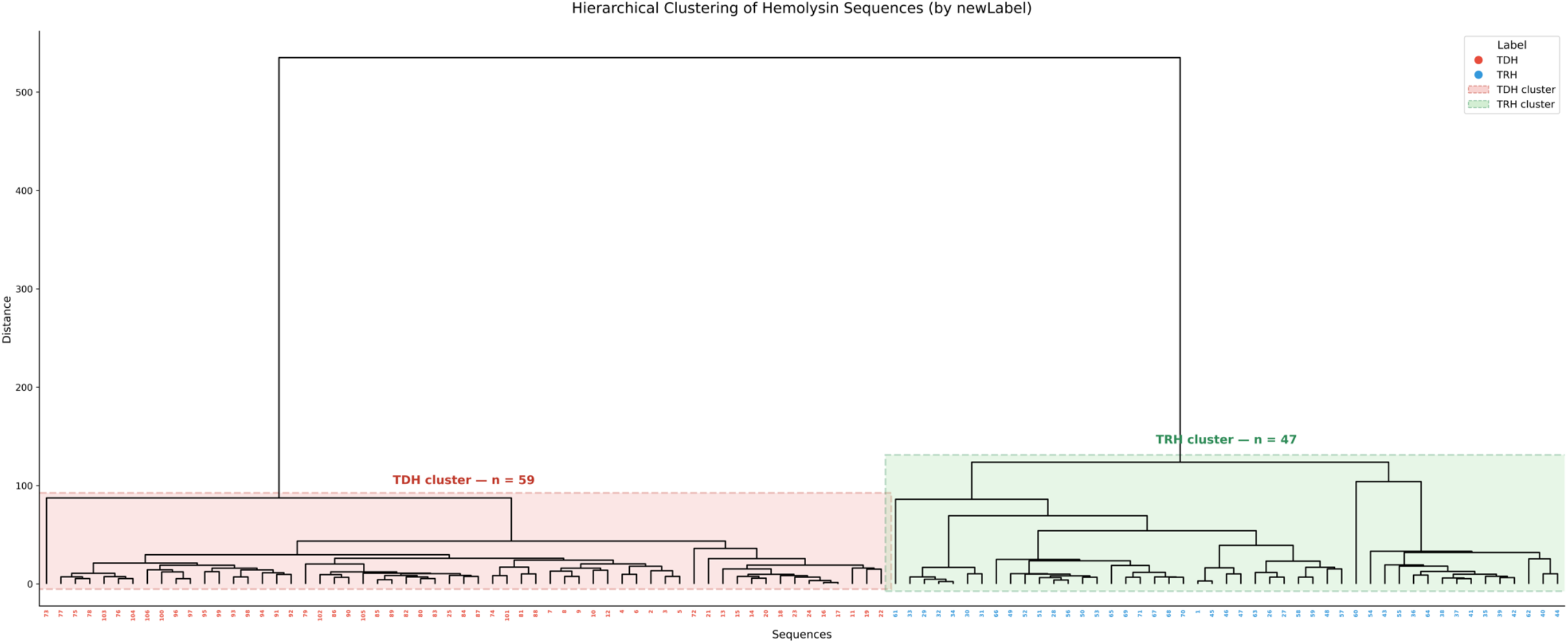
Dendrogram of *Vp* Haemolysin based on similarity at the amino acid sequence level.

### TDH and TRH exhibited similar global representation independently by forming distinct clusters revealed by ESM2

The inferred functional embedding of TDH and TRH were pairwise compared based on cosine similarity. The lowest cosine similarity was more than 0.957, which indicated that TDH and TRH overall showed similar functional properties. Noticeably, the distinct and separated clustering pattern between TDH and TRH was observed on UMAP results regardless of high global representation similarity (**Fig 4**.). The separated clusters between TDH and TRH resonated the previous statement may support the finding that TDH and TRH form distinct structural clusters despite maintaining a high global functional similarity. In addition, the intra-class sub-clustering pattern differed between the TDH and TRH groups. The TDH embeddings exhibited a relatively tight clusters, whereas the TRH embeddings exhibit a wider dispersion with a distinct gap in the embedding space, which implied that TDH had lower variability in sequence-encoded features than TRH.

**Figure 4.**
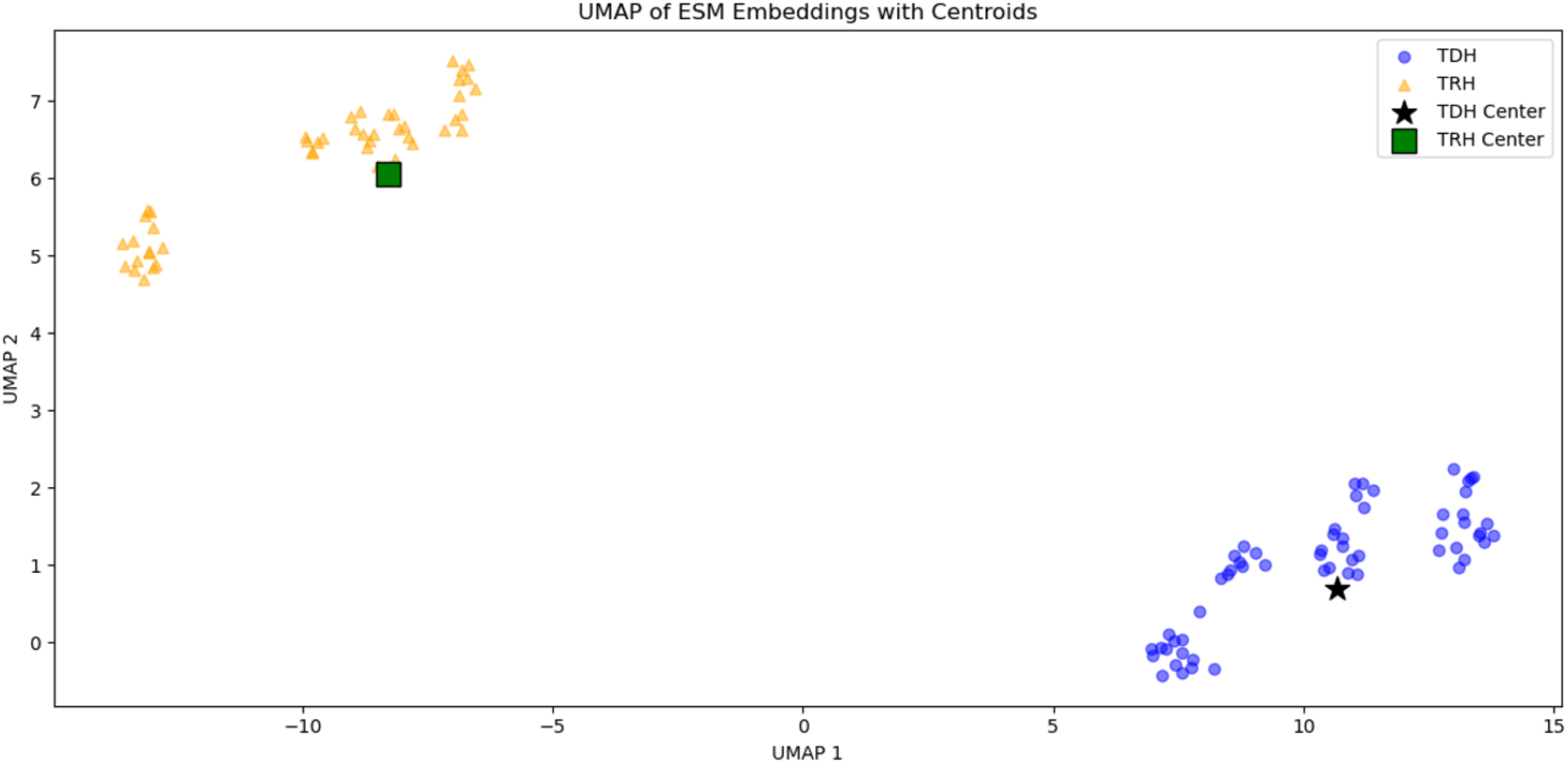
UMAP of ESM-derived protein representations for thermostable direct haemolysin (TDH) and TDH-related haemolysin (TRH) from *Vp*.

### Consistent yet secondary structure diverse haemolysin protein structures were inferred by AlphaFold

Folding structures of 106 amino acids for TDH and TRH were further inferred using AlphaFold2 (**S Fig 6.** and **S Fig 7.**). AlphaFold2 predictions for all sequences showed consistently high structural quality, with mean pLDDT values ranging from 80–83, which indicated reliable backbone confidence and pTM scores of ∼0.74–0.78 supporting correct global folding. Predicted alignment error remains stable across all groups and mean off-diagonal PAE (∼10–11.7 Å) suggesting moderate but expected inter-domain flexibility rather than structural uncertainty. Both TDH and TRH exhibit extended α-helical regions coupled with compact core domains, suggesting conserved structural motifs that may underpin their common biological functions in virulence. Essentially, AlphaFold results reveals previous conserved regions identified by sequence alignment were consistent with the beta-sheet structure of TDH/TRH (S Fig 8.). The extended α-helices in both structures are commonly self-highlighted, suggesting important interactions with either membranes or binding partners. Within each haemolysin clusters, two subclusters were shown for TRH and four subclusters were shown for TDH. Energy evaluation using the Rosetta scoring function was further implemented to validate the structure prediction of AlphaFold, which enabled evaluation of the energetic plausibility of predicted folding structures and facilitated comparative analysis of stability across TDH and TRH variants. TRH generally had lower Rosetta energy unit (REU) compared with TDH (**S Fig 11**.), which indicated more stable internal energy of a static fold. and thermodynamically favorable protein folding structure. K-means clustering was applied on TDH and TRH ESM2 embeddings separately and the cartoon protein folding structure of TDH/TRH closest to the cluster centroid were visualized (**S Fig 9. and S Fig 10**.). The cluster central representative folding structure of TRH all showed alpha helix in the tail region, whereas TDH counterparts showed coil structure and beta-sheet, which indicated that compact and energetically optimized local conformation from the prevalence of alpha-helical motifs in the TRH tail region may be associated with its lower REU. Lipid membrane affinity of TDH and TRH were further simulated based on structural inference by AlphaFold2. Based on the membrane affinity simulation results (**S Fig 12**.), no significant difference between the membrane affinity of TDH and TRH (i.e. mean COM distance, min COM distance, and mean contact beads), although TDH showed slightly higher mean contact beads and lower min COM distance than TRH. Although TDH and TRH exhibit distinct patterns in structural folding, membrane affinity (indicative of initial stage of haemolysin-membrane interaction) is not functional consequence of structural difference. Instead, these observed structural differences may be associated with the downstream haemolysin-membrane interactions such as pore formation and ion flux induction (Takahashi et al., 2000; Verma & Chattopadhyay, 2021). Raghunath (2015) summarized that TDH forms approximately 2-nm membrane pores that permit the passage of water and ions, while TRH, similar to TDH, activates Cl⁻ channels and thereby alters intestinal ion flux, contributing to diarrhea.

### Combined functional and structural representation learning revealed the critical accumulated point mutations from TDH to TRH

Nishibuchi and Kaper (1995) stated that TDH and TRH are likely from the same ancestor and diverging because of accumulated point mutations throughout time. To further reveal how point mutations and accumulation of them lead to functional divergence between TDH and TRH, gradient boosting trees were utilized to select position and amino acid residue with importance contributing to the differentiation between TDH and TRH. The cosine similarity landscape toward TDH and TRH central embeddings was shown in **Fig 5a.,** of which the overall clustering pattern was consistent with UMAP results. **Fig 5b**. illustrated a stepwise TRH-directed mutation path, where point mutations identified by a gradient boosting tree progressively shift the ESM2 embedding, and their accumulated effect formed a continuous trajectory from TDH toward the TRH region in cosine similarity space. A total of 23 steps were identified on the cleaned path from TDH to TRH, which contained a collection of single point mutation steps and steps involving accumulated point mutations. The identified point mutations are distributed across the sequence rather than concentrated at specific sites and resulted sequence of each step was structurally predicted using AlphaFold (**Table 1 and S Fig 13**). The biological shift from TDH toward TRH is a sequence mutation in a full spectrum of 189-position amino acid pattern, which suggests that correlated changes among distinct protein regions may be important. The folding structure of collection of mutated amino acid sequence was further inferred again using AlphaFold, and results suggested that as the mutations identified by gradient boosting tree continue to accumulate, the tail region of haemolysin began to form alpha-helix.

**Figure 5.**
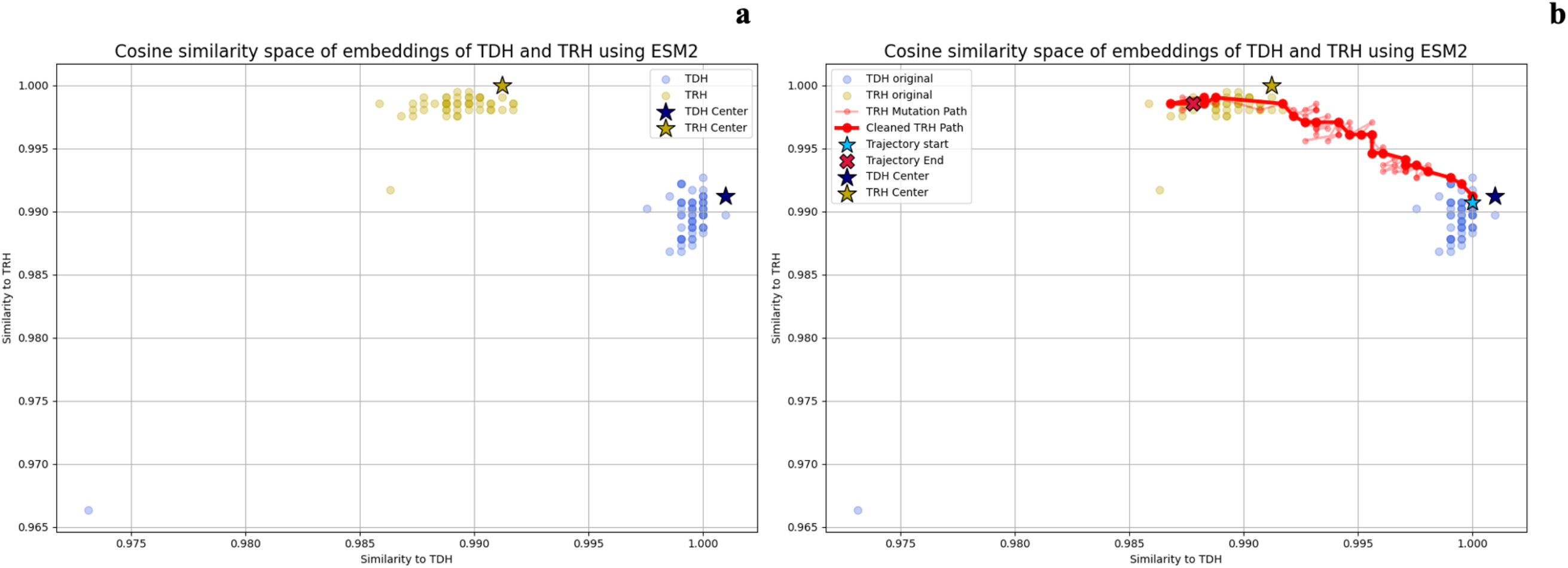
Cosine similarity–based embedding space of ESM-derived protein representations for thermostable direct haemolysin (TDH) and TDH-related haemolysin (TRH) from *Vp*.

**Table 1:**
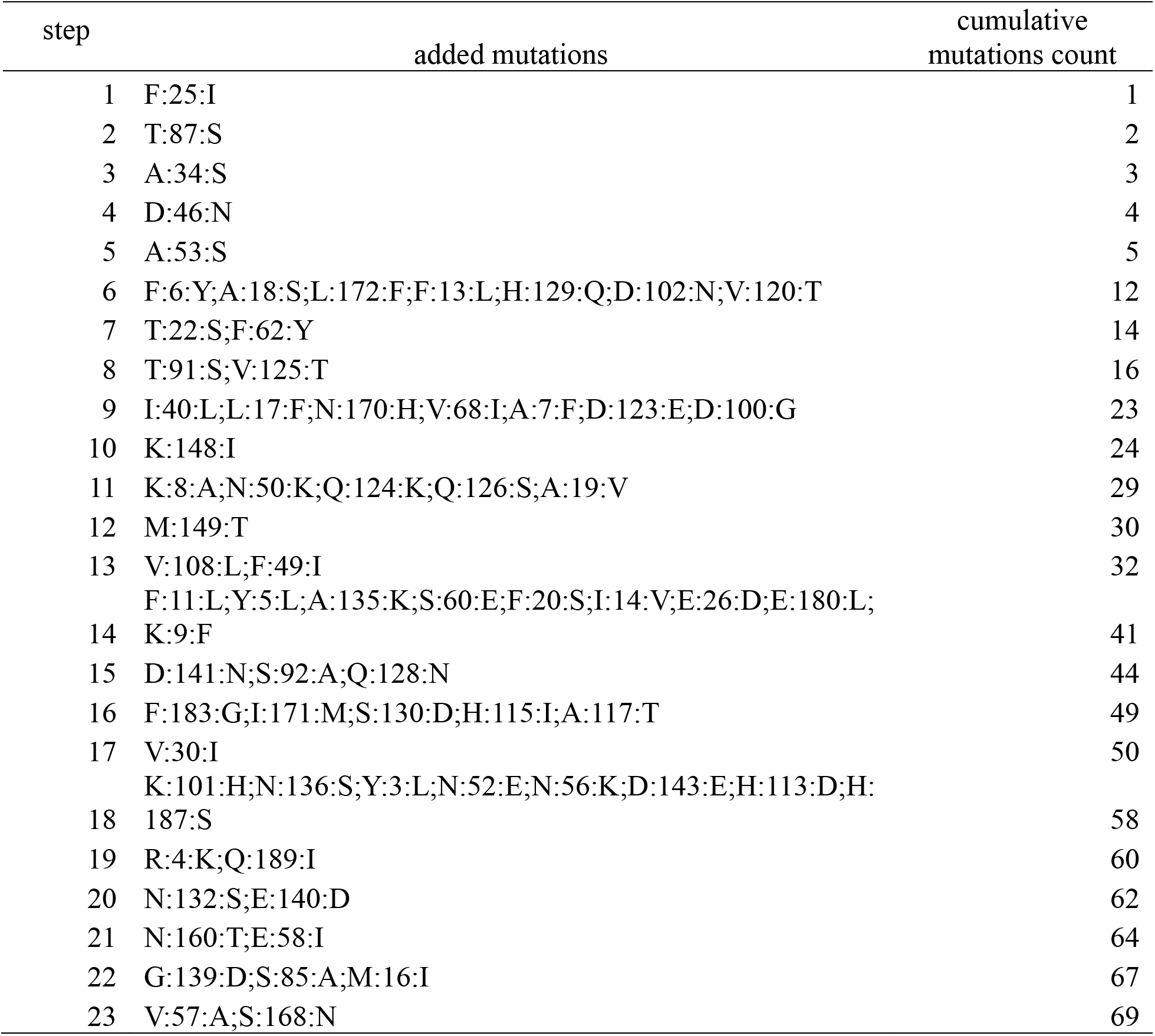
Identified accumulated point mutations contributing to TDH-to-TRH divergence.

## Discussions

Omics-driven scientific discovery and innovative application have been previously conducted on *Vp* at genomics and transcriptomics levels (Z. Liu et al., 2025; Z. Liu et al., 2023). Targeting *Vp* haemolysin characterization, this study extended such efforts to the proteomic levels. The integrative analysis of TDH and TRH in this study consists of amino acid sequence-level bioinformatics analyses and protein structural and functional representation learning discovery, which aimed to describe a full spectrum of their genetic variations, functional properties, and biophysical implications. At the amino acid sequence level, the bioinformatics analytics focused on the identification of key mutations and sequence conservation that may contribute to the main differentiation between TDH and TRH. At the protein functional and structural levels, advanced deep learning–based protein structure prediction (AlphaFold) and protein language models (ESM2) were used to obtain the protein folding structures and infer the functional similarity between TDH and TRH, respectively. Given the leading effects on *Vp* pathogenicity, TDH and TRH serve as the major virulence factors and their deriving *tdh* and *trh* genes have been widely used as reliable indicator in detection methods (Liao et al., 2017; Nilsson & Turner, 2016).

Traditionally, deriving structural representations and functional insights for proteins has been both time-consuming and resource-intensive, which has been a major bottleneck that prevents large-scale analysis of *Vp* haemolysin (Akdel et al., 2022; Huang et al., 2023). To our best knowledge, this study is the first scalable examination leveraging representation learning to efficiently characterize the structural and functional landscape of *Vp* haemolysin.

A substantial amount of point mutations across 189 amino acid bases were observed based on the increasing number of amino acid sequences of *Vp* haemolysin deposited into public databases.

Previous consensus was that the amino acid sequence similarity between TDH and TRH was approximately 67-70% (Gutierrez West et al., 2013). In contrast, the similarity analysis between TDH and TRH at amino acid sequence level (**Fig. 4 and Fig. 5**), showed relatively lower identity percentage similarity (56.1% to 67.4%) between TDH and TRH at amino acid levels. Such discrepancy may be attributed to several confounding factors, for instance sampling bias in earlier studies, where *Vp* strains were derived from a limited geographic region or isolation types and a limited number and types of strains (Siddique et al., 2021; Yanagihara et al., 2010), whereas the present work provided a systematic and comprehensive analysis of all currently available TDH and TRH sequences, capturing a much wider spectrum of genetic divergence.

When viewed solely at the amino acid sequence level, such observed haemolysin amino acid sequence variability appeared to be biological context deficient, motivating the use of representation learning to uncover latent functional and structural patterns behind monotonous and even chaotic point mutation patterns.

An unprecedented multi-dimensional understanding of TDH and TRH was achieved by integrating insights from amino acid sequence, functional representation, and structural folding prediction. Protein language model driven ESM2 embeddings of *Vp* haemolysin serve as promising tool for functional comparison between TDH and TRH. Regardless the substantial divergence between TDH and TRH at the amino acid sequence level, TDH and TRH exhibited highly similar cosine similarity values in the embedding space, while still forming well-separated clusters, which suggested that TDH and TRH retain conserved higher-order but nuanced functional and structural features. Based on the AlphaFold protein folding inference results, membrane affinity of all 106 TDH and TRH were further investigated. Lipid-protein simulations are critical for *Vp* haemolysin, reported as pore-forming toxin target cellular membrane in previous literature, which enable the systematic, high-throughput exploration of biologically complex interaction between protein and cellular membranes (Wassenaar et al., 2015). No significant difference (P>0.05) was observed on simulated membrane affinity evaluation metrics between TDH and TRH, which mirrored embedding space observations and suggested that shared membrane-interacting functionality of TDH and TRH is conserved despite sequence-level divergence. It is noteworthy that TDH and TRH are in tetramer format when actively attacking host cells. Further investigation of membrane affinity of TDH and TRH in tetramer is warranted. Finally, key amino acid positions across the 189-residue TDH and TRH were identified using gradient boosting trees, and these positions were consistently supported by both functional and structural representation learning. The function change from point mutations may include but not limited to nuance in membrane pore forming strategy (direct pore formation by TDH and Cl⁻ channel-mediated ion flux by TRH) and structural difference in haemolysin tail regions folding (alpha-helix on TRH tail) based on insights retrieved from this study.

## Conclusion

This study leveraged bioinformatics analytics at the amino acid sequence level and representation learning at the protein functional and structural levels to reveal, for the first time, the multi-dimensional divergence between TDH and TRH. Outcomes revealed distributed, accumulated point mutations across the full-length protein. Despite pronounced sequence-level divergence, both toxins retain highly similar functionally embeddings and share core virulence mechanisms. By integrating sequence analytics, protein language model embeddings, and structure-based simulations, this study also indicated the potential toxin function change lead by mutation, such as the pore-forming mechanisms (direct pore formation by TDH versus Cl⁻ channel activation by TRH) and hemolysin tail folding (an α-helix in the TRH tail). Insights from this study reinforce the rationale using *Vp* strain harboring *tdh* and *trh* genes in experimental design. The outcomes from this study have provided complementary information on TDH and TRH as well-known virulence factors, thereby enhancing understanding of microbial pathogenesis of *Vp*. The representation learning driven methods demonstrate the potential to be used for studying a wider range of virulence factors of *Vp* and toxins of other pathogenic foodborne bacteria.

## Data Availability

The data and code used for analysis are available in the GitHub repository at https://github.com/jlk666/tdh_trh_Vp,

## Acknowledgement

We sincerely thank Dr. Linda Harris and Dr. Crystal (Xiang), Yang’s insights and revision suggestions on the manuscript.

---

**Supplementary Figure 1.**
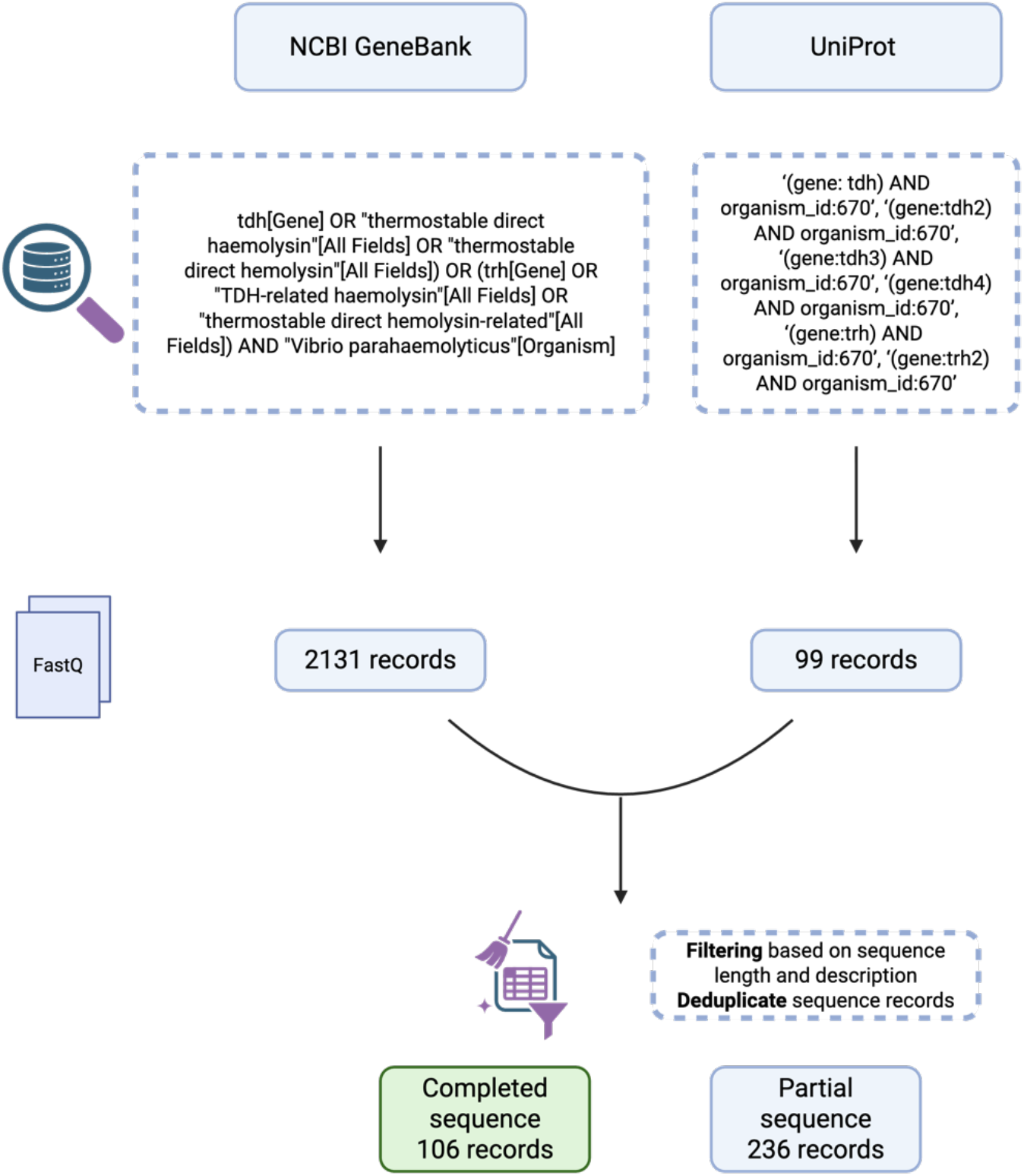
Overview of TDH/TRH sequence collections from public databases and data wrangling pipeline used in this study.

**Supplementary Figure 2:**
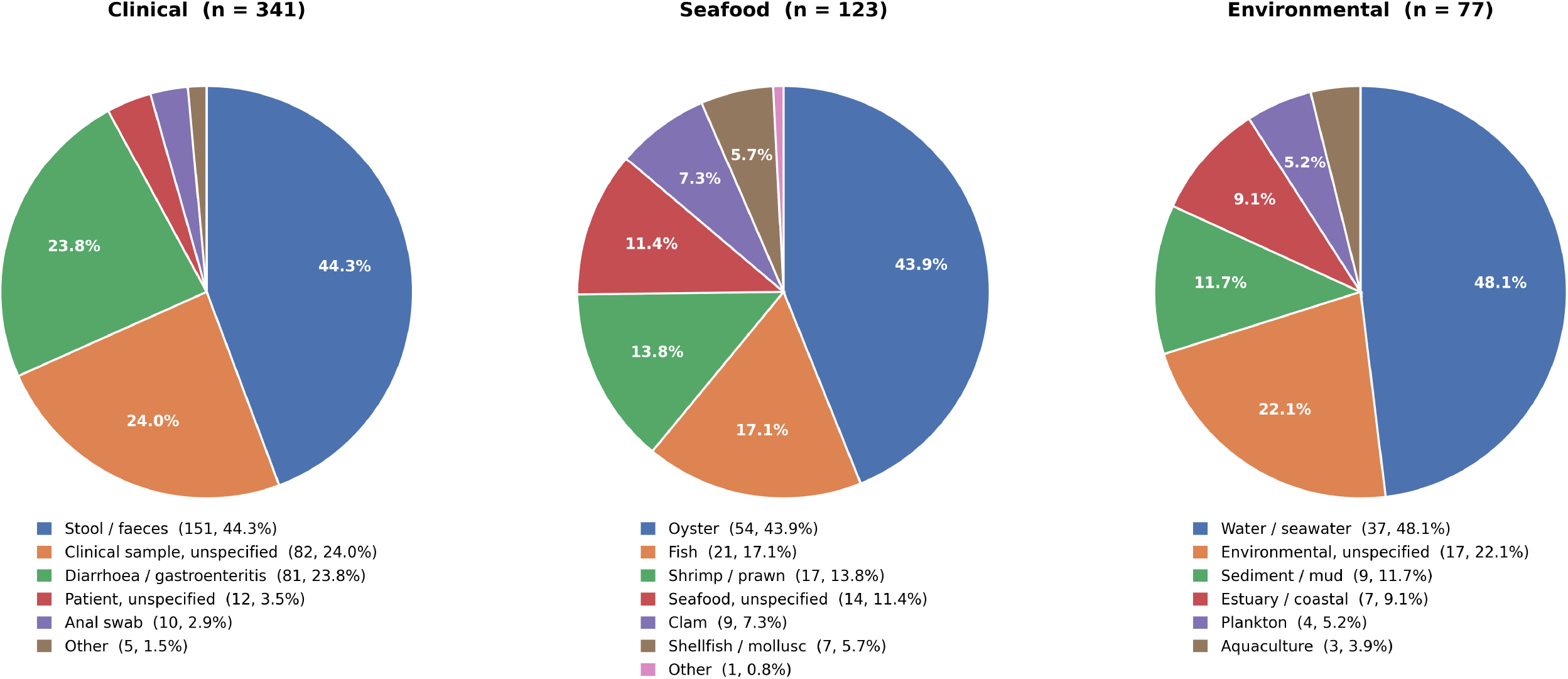
Pie charts showing distribution of strain isolation source of *Vp* strain harboring TDH/TRH: clinical, seafood and isolation based on available metadata information of collected NCBI data.

**Supplementary Figure 3:**
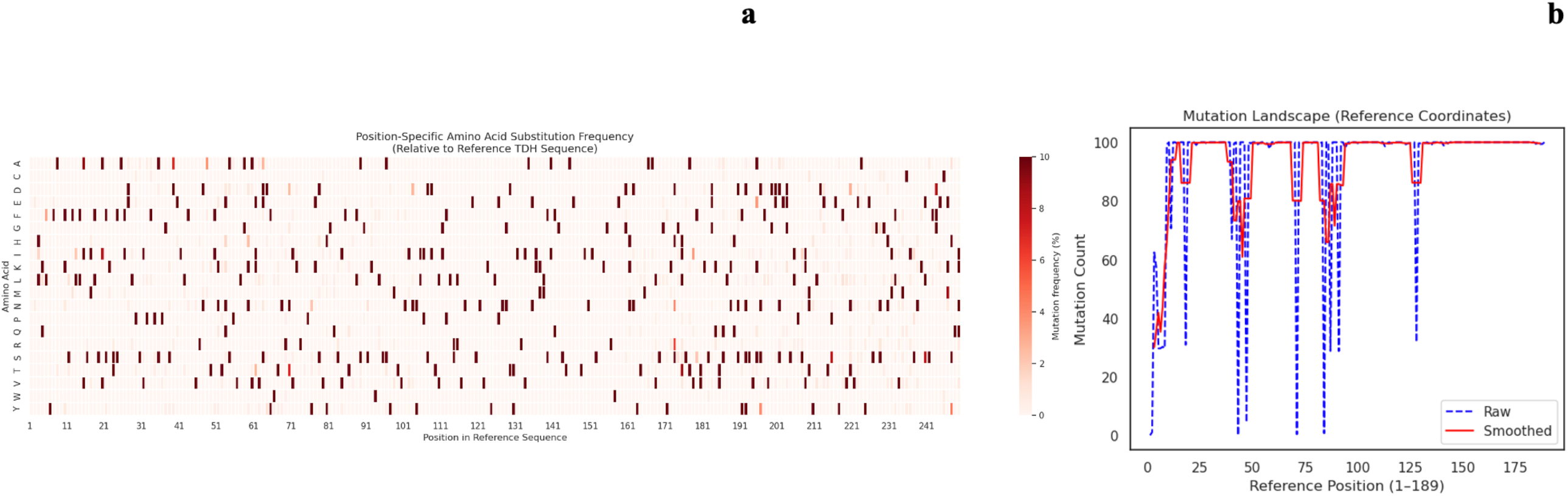
Comparative analysis of position-specific mutation frequencies against reference sequence using aligned complete and incomplete of *Vp* haemolysin amino acid sequence (a) and mutation landscape (b).

**Supplementary Figure 4:**
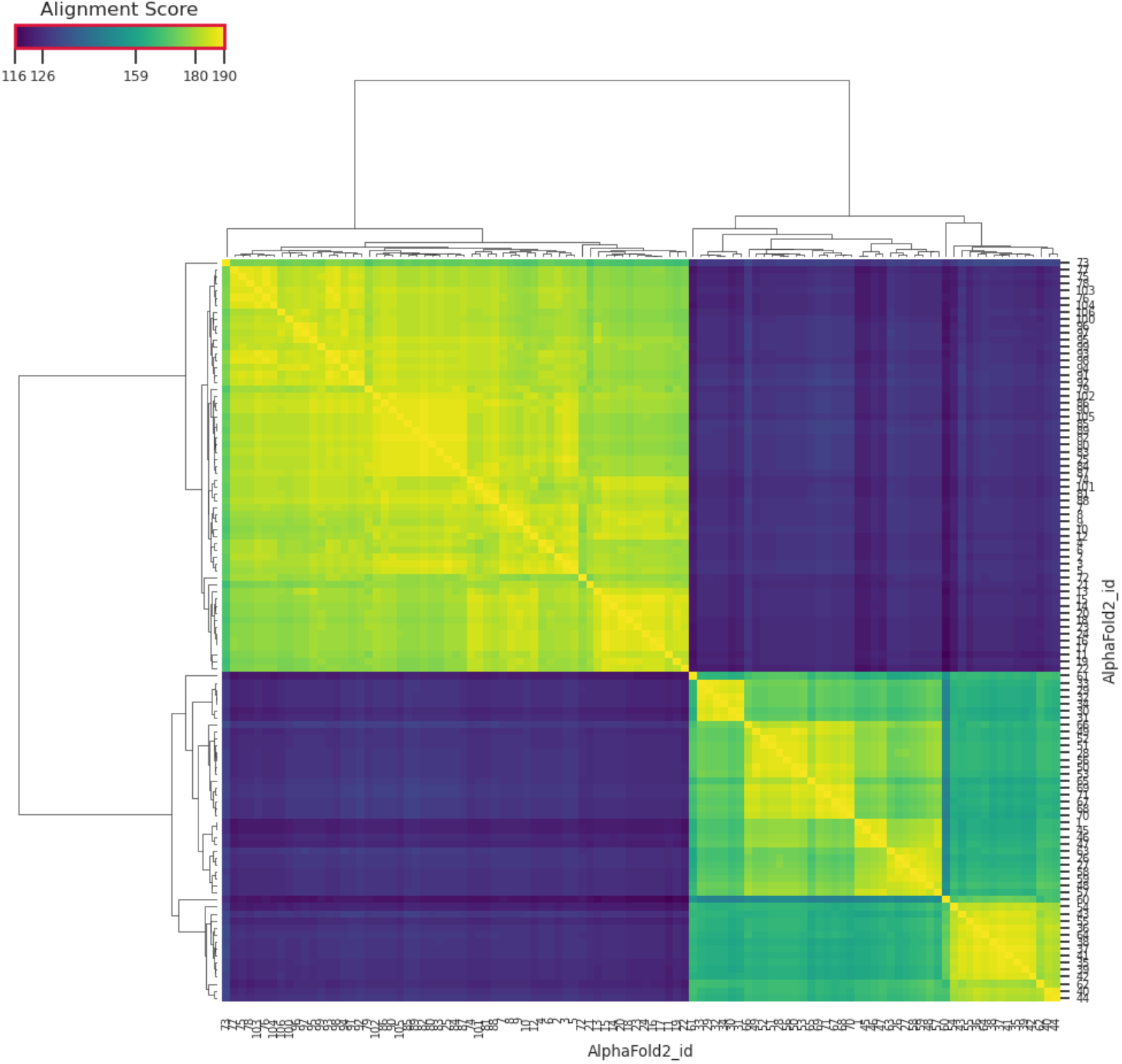
Hierarchical Clustering and Similarity Heatmap of *Vp* Haemolysin.

**Supplementary Figure 5:**
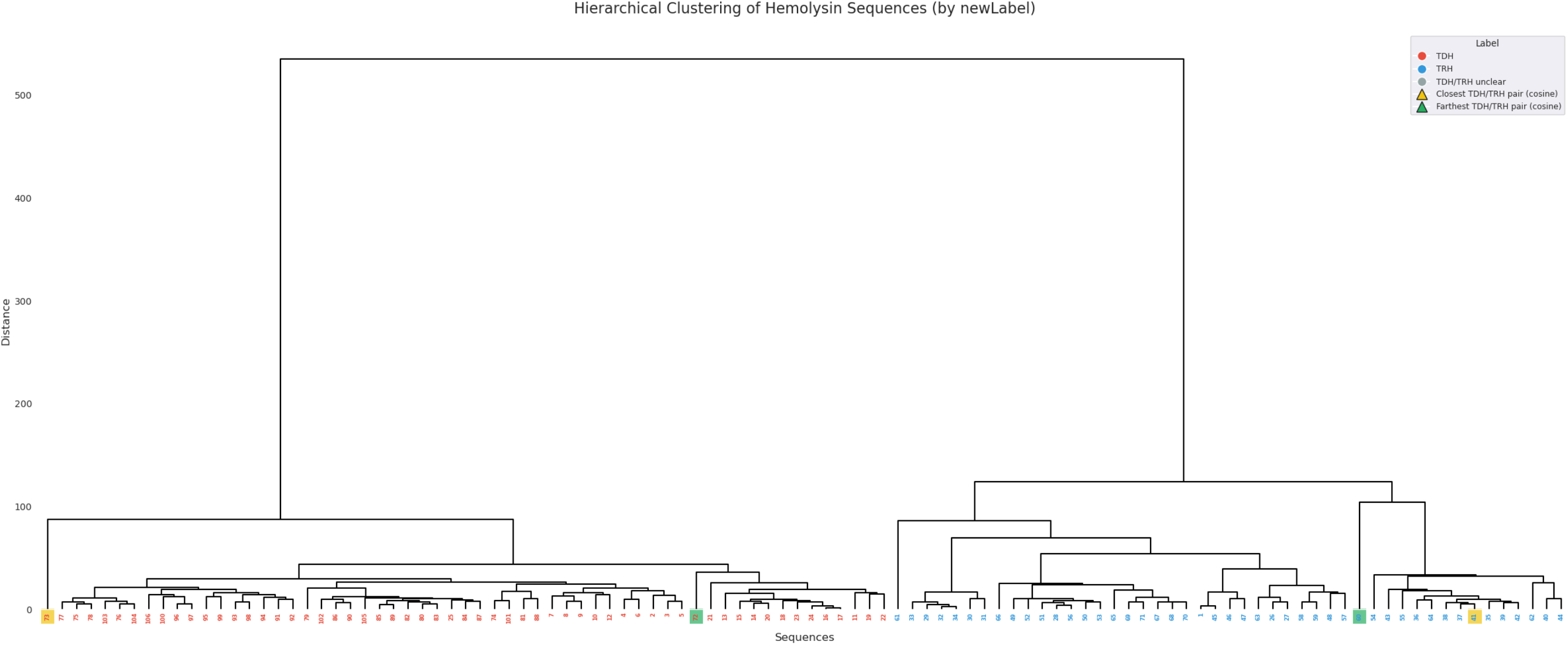
Complete TDH and TRH sequences were highlighted based on percent identity distance (Green and yellow show the lowest and highest similarity respectively).

**Supplementary Figure 6:**
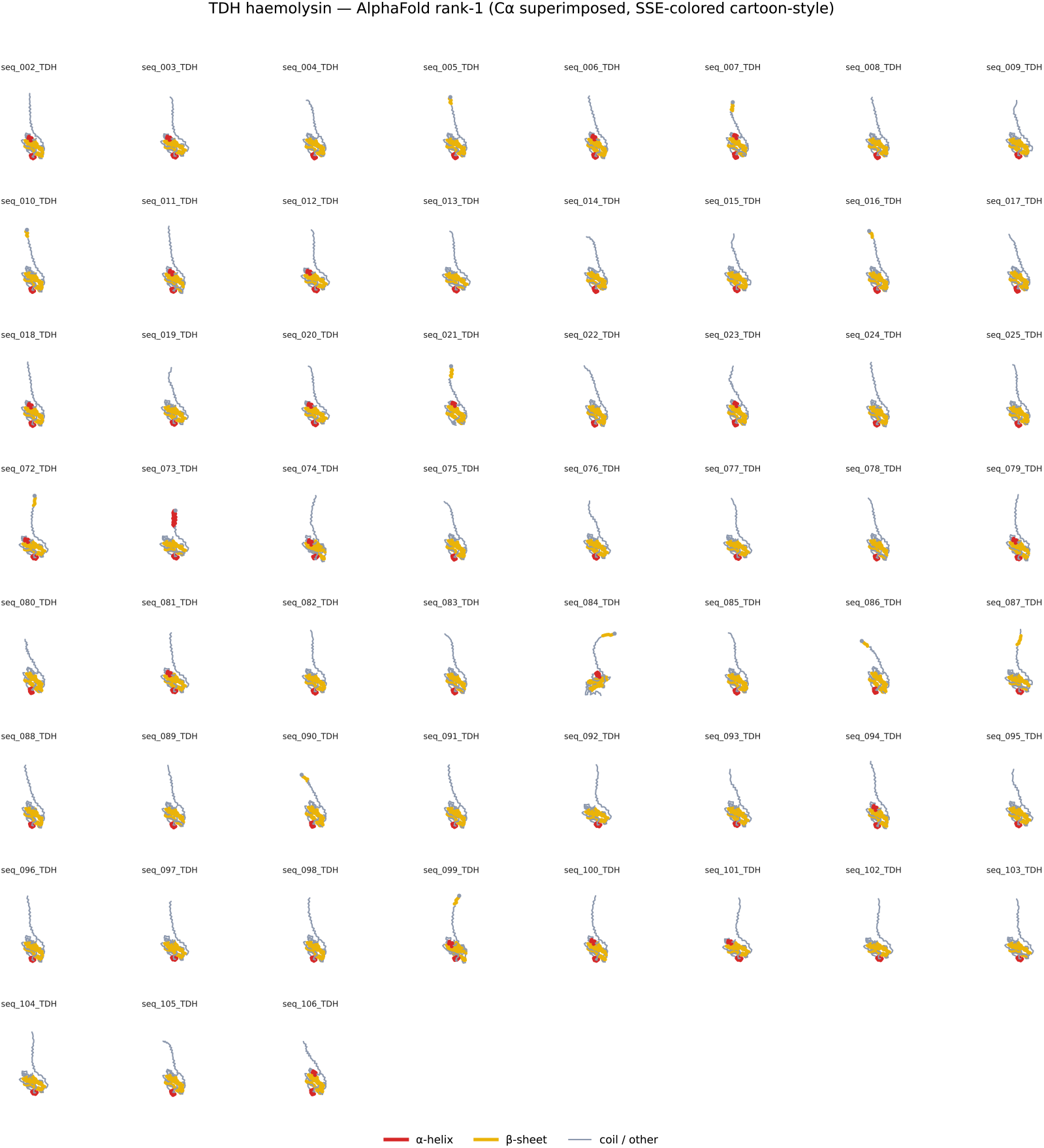
AlphaFold-predicted structures of TDH

**Supplementary Figure 7:**
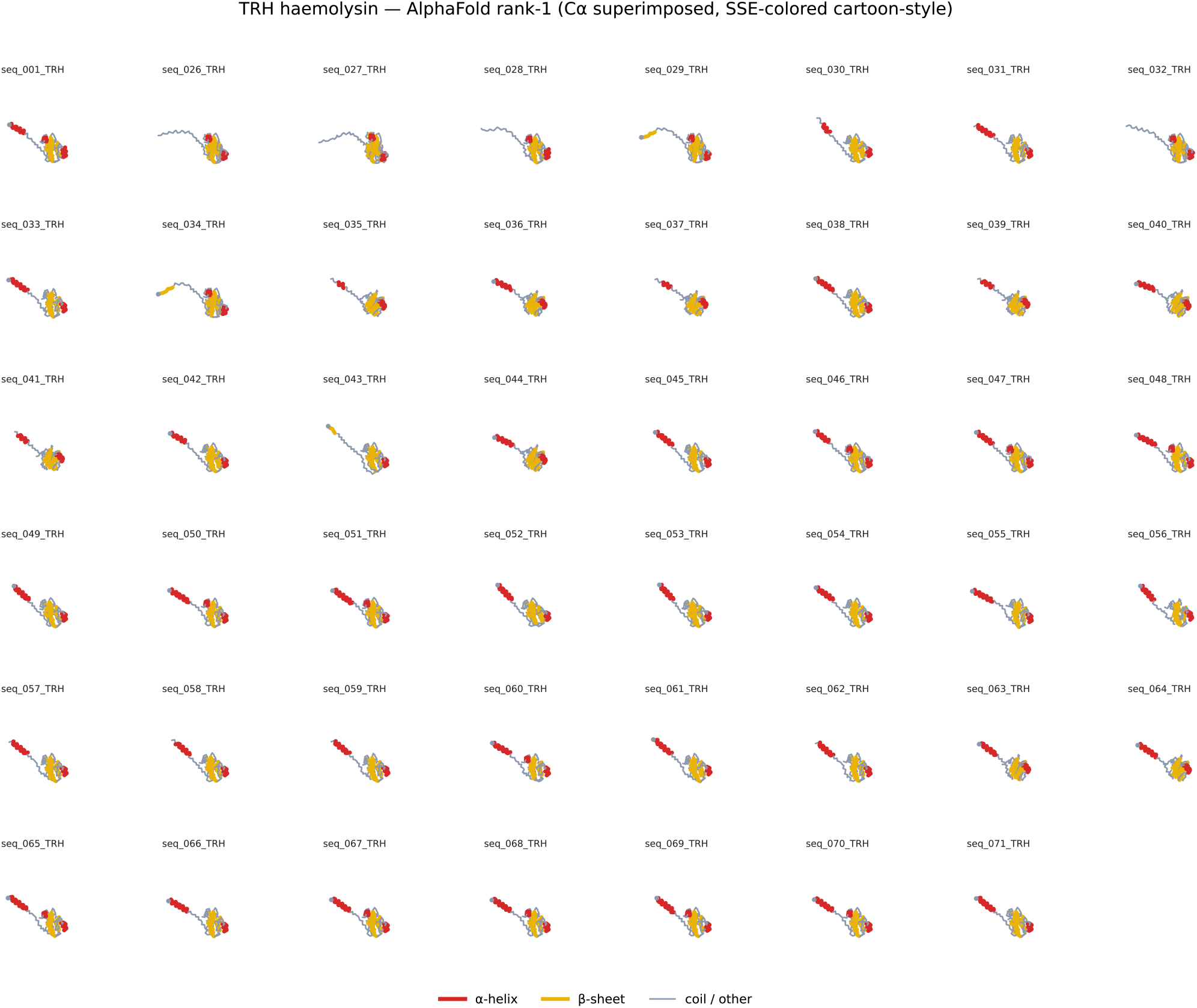
AlphaFold-predicted structures of TRH

**Supplementary Figure 8:**
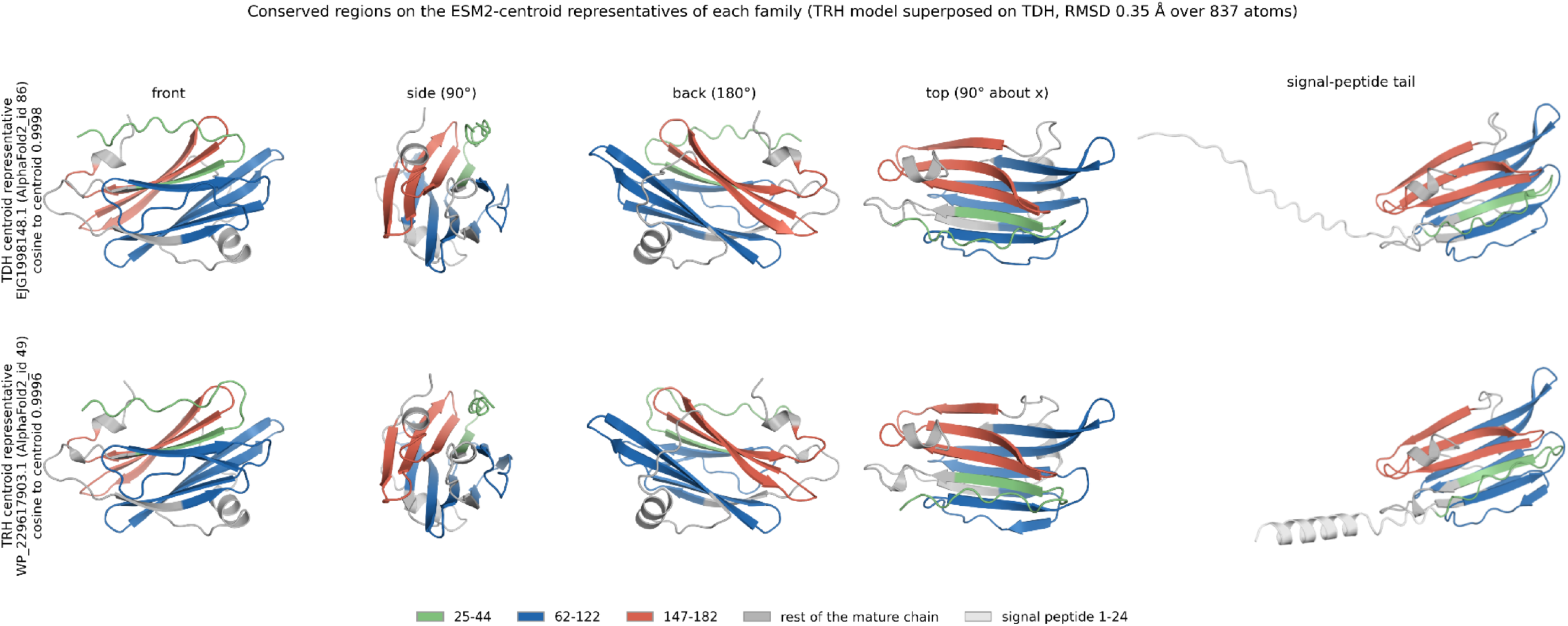
Conserved regions mapped onto ESM2-centroid representative structures of TDH and TRH. AlphaFold2- predicted structures of TDH (top) and TRH (bottom) are shown from front, side, back, top, and signal-peptide-tail views. Conserved regions at positions 25–44, 62–122, and 147–182 are highlighted in green, blue, and red, respectively.

**Supplementary Figure 9:**
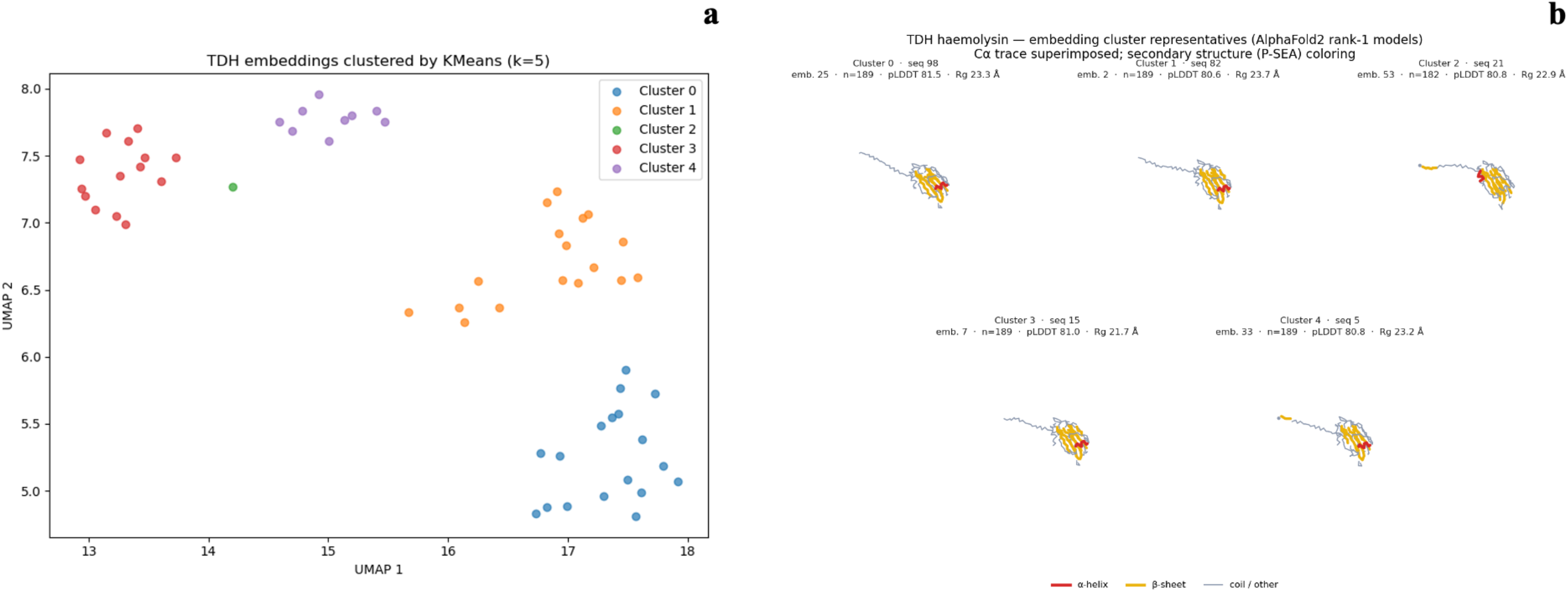
ESM2-embedding of TDH were further clustering using K-mean (k = 3) (**a**). The actual TDH protein structures from AlphaFold representing each cluster centers were visualized with alpha-helix, beta-sheet, and coil colored in red, yellow and grey collectively (**b**).

**Supplementary Figure 10:**
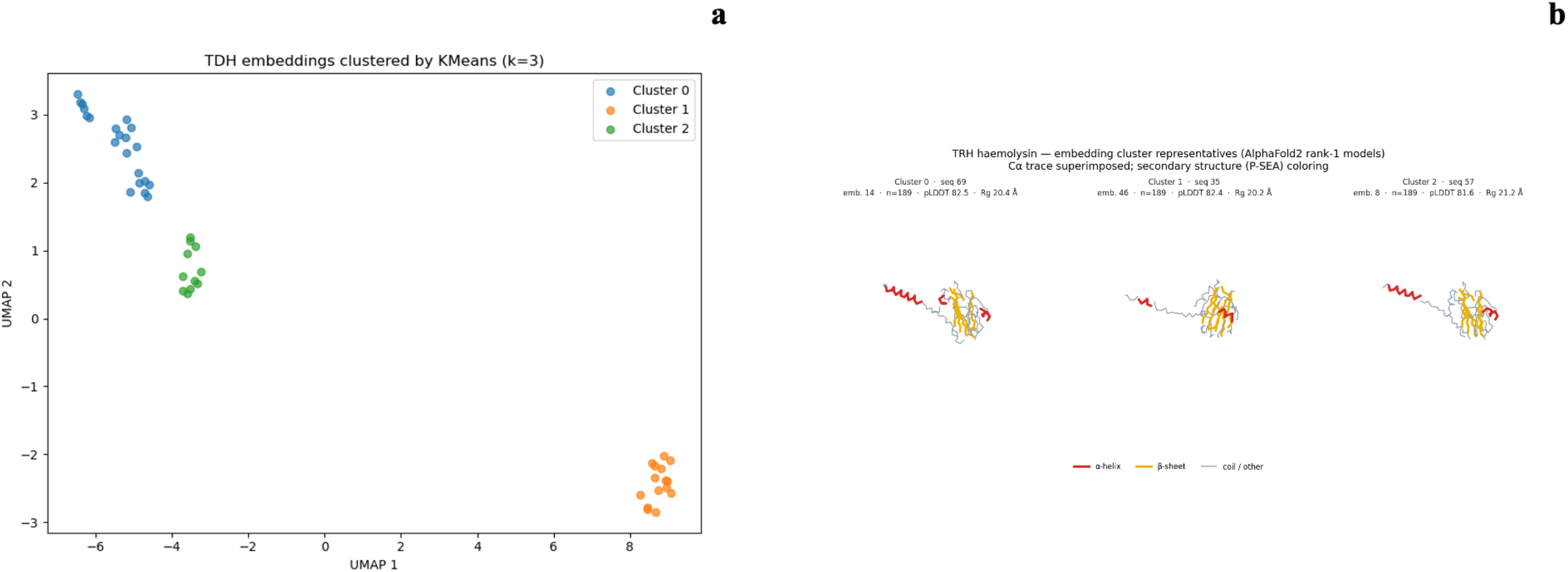
ESM2-embedding of TRH were further clustering using K-mean (k = 3) (**a**). The actual protein structures from AlphaFold representing each cluster centers were visualized with alpha-helix, beta-sheet, and coil colored in red, yellow and grey collectively (**b**).

**Supplementary Figure 11:**
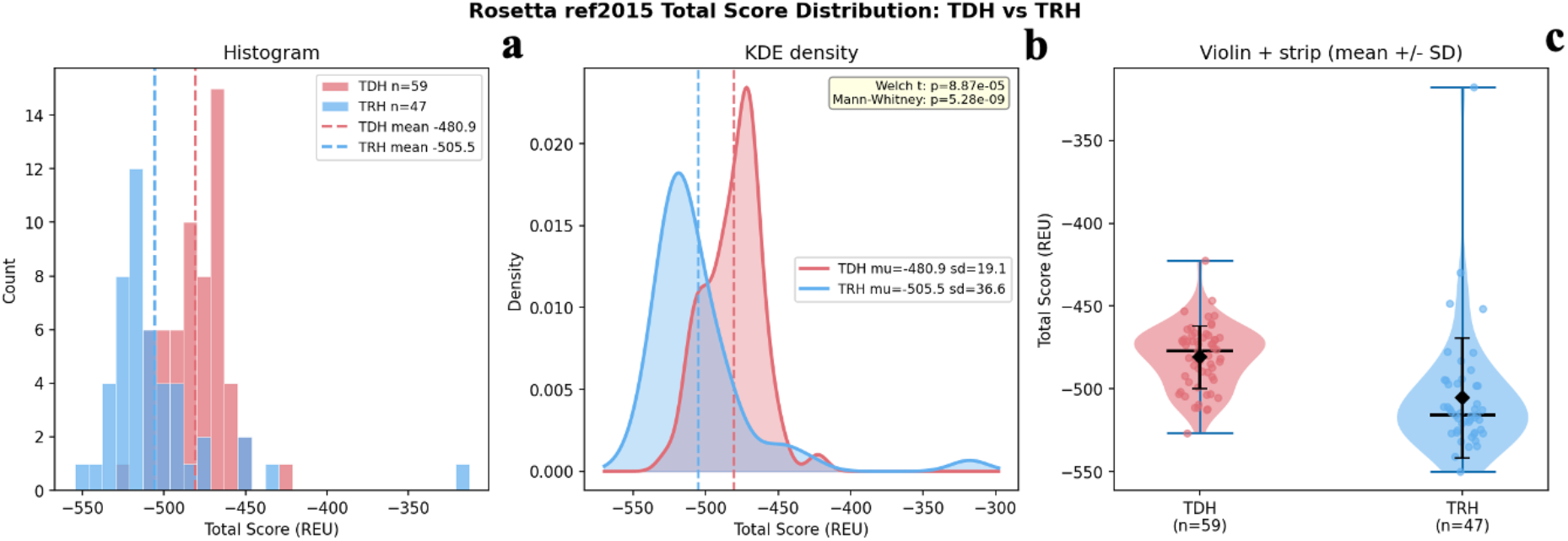
Histogram of total scores (REU) for TDH (red, n = 59) and TRH (blue, n = 47); dashed lines indicate means (**a**). KDE plots showing distribution differences; p-values from Welch’s t-test and Mann–Whitney U test are indicated (**b**). Violin plots with individual points and mean ± SD for TDH and TRH (**c**).

**Supplementary Figure 12:**
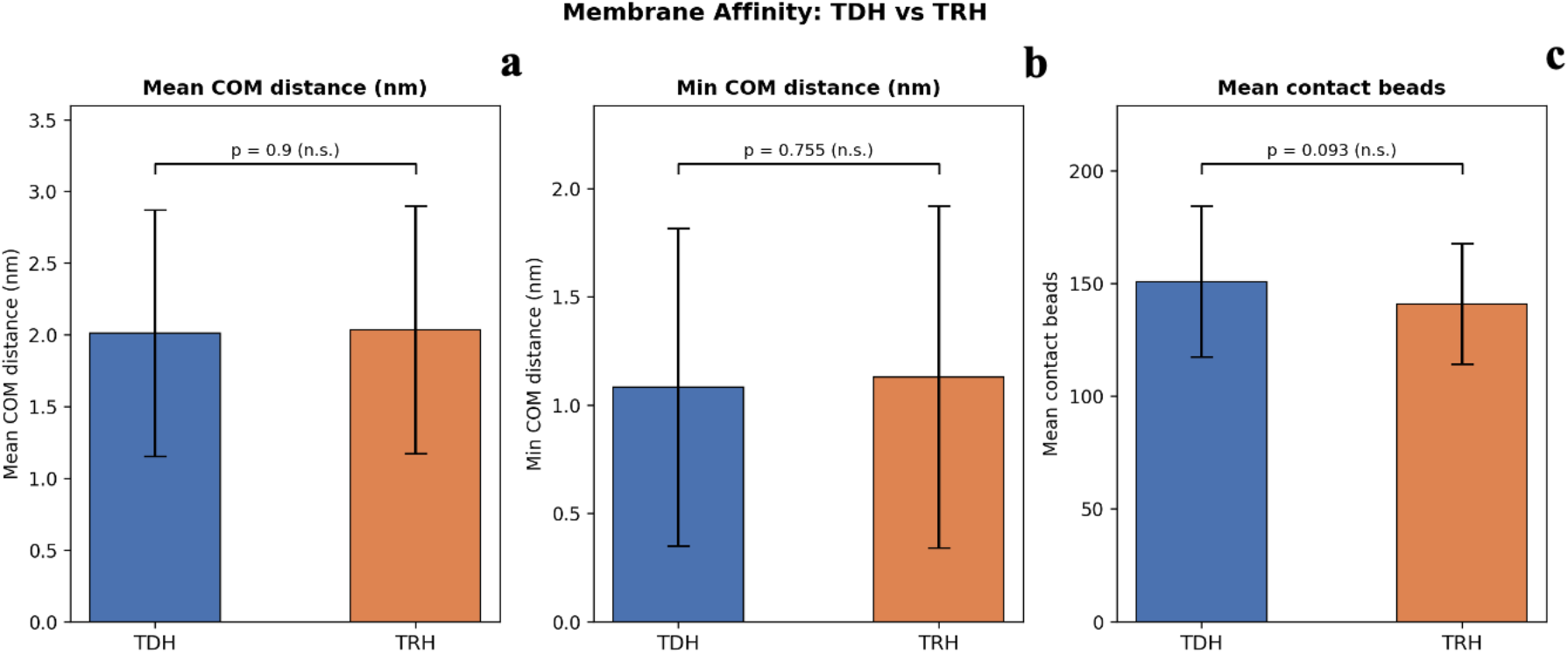
Membrane affinity comparison between TDH and TRH. Mean COM distance to the membrane (nm) for TDH and TRH (a). Minimum COM distance to the membrane (nm) (b). Mean number of contact beads (c). P-values are indicated; all comparisons are not significant.

**Supplementary Figure 13:**
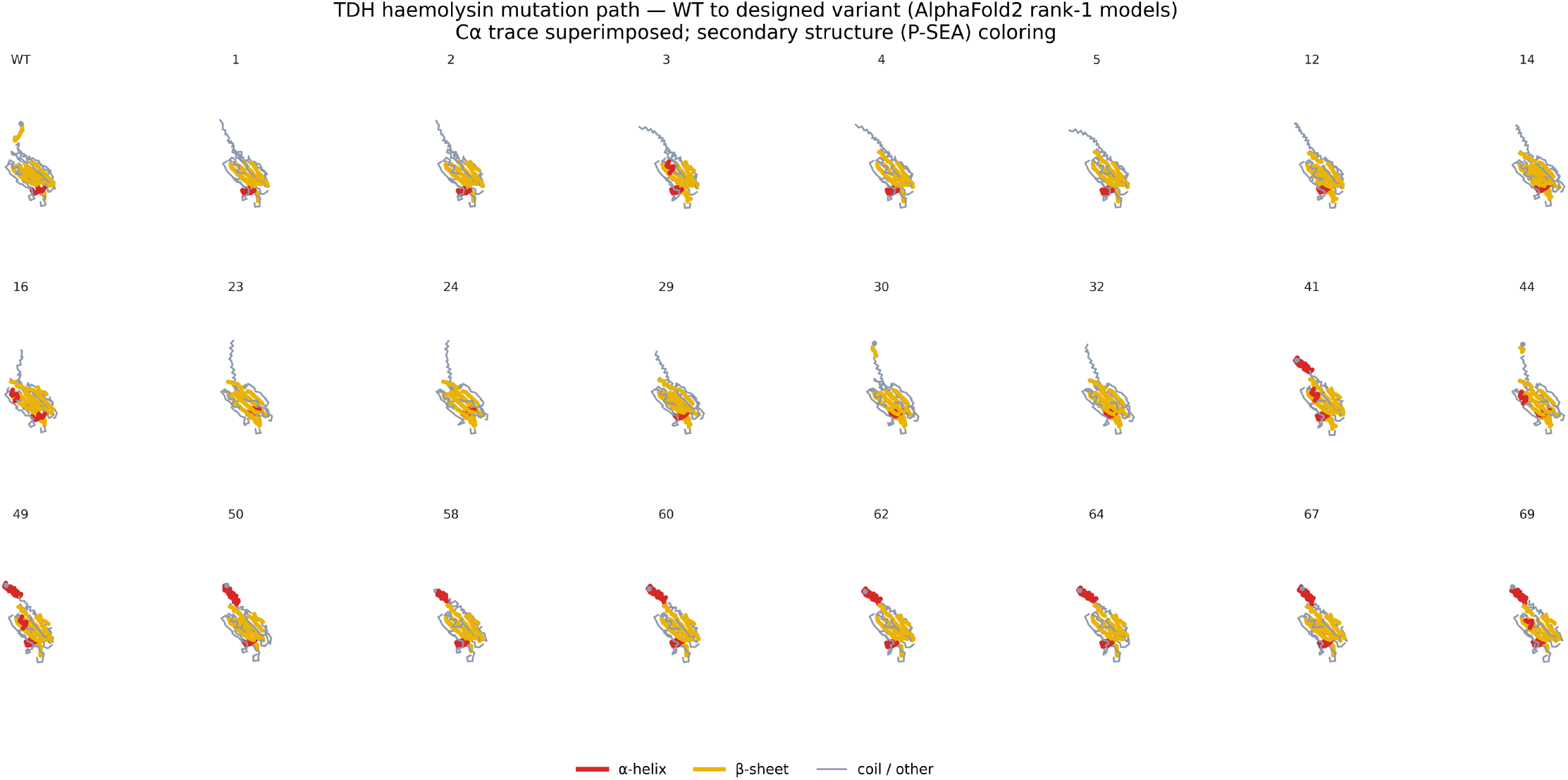
AlphaFold-predicted structures of along the TDH-to-TRH with accumulated point mutations.

